# simGWAS: a fast method for simulation of large scale case-control GWAS summarystatistics

**DOI:** 10.1101/313023

**Authors:** Mary D. Fortune, Chris Wallace

## Abstract

**Motivation:** Methods for analysis of GWAS summary statistics have encouraged data sharing and democratised the analysis of different diseases. Ideal validation for such methods is application to simulated data, where some “truth” is known. As GWAS increase in size, so does the computational complexity of such evaluations; standard practice repeatedly simulates and analyses genotype data for all individuals in an example study.

**Results:** We have developed a novel method based on an alternative approach, directly simulating GWAS summary data, without individual data as an intermediate step. We mathematically derive the expected statistics for any set of causal variants and their effect sizes, conditional upon control haplotype frequencies (available from public reference datasets). Simulation of GWAS summary output can be conducted independently of sample size by simulating random variates about these expected values. Across a range of scenarios, our method, produces very similar output to that from simulating individual genotypes with a substantial gain in speed even for modest sample sizes. Fast simulation of GWAS summary statistics will enable more complete and rapid evaluation of summary statistic methods as well as opening new potential avenues of research in fine mapping and gene set enrichment analysis.

**Availability and Implementation:** Our method is available under a GPL license as an R package from http://github.com/chr1swallace/simGWAS

**Supplementary Information:** Supplementary Information is appended.

## Background

The genome wide association study design is now more than a decade old (Visscher et al., 2017), and the size of GWAS cohorts has continued to grow, from 1000s to, now, 1,000,000s of individuals. Given the competing demands of open science and privacy concerns (P3G Consortium et al., 2009), it has become standard to share data in the form of summary statistics (allelic effect sizes and standard errors, or simply p values) more readily than the full genotype data. A wealth of methods have been developed to operate directly on the summary statistics, from fine mapping of genetic causal variants (e.g. PAINTOR (Kichaev et al., 2014), CAVIARBF (Chen et al., 2015) and JAM (Newcombe et al., 2016)) to (co-)heritability estimation (Bulik-Sullivan et al., 2015) and integration of GWAS results from different traits (Giambartolomei et al., 2014; Zhu et al., 2016). Summary data methods are often derived through approximating a multivariate linear regression likelihood by incorporating information about correlation structures (linkage disequilibrium, LD) from reference populations. However, one must adopt a logistic regression approach to correctly model risk on the log odds scale when analysing GWAS of binary traits (including case-control data). Summary statistic methods which have been originally derived for linear regression cannot do this and the impact of the linearity assumption on their conclusions if applied to case-control data has not been investigated in depth.

As Biobank-sized datasets come to fruition, such summary statistic methods are likely to become even more important, since, for such large numbers of samples, operating on the complete genotype data matrices for efforts such as Bayesian fine mapping of causal variants is computationally prohibitive. Indeed, GWAS summary statistics for multiple traits from UK-Biobank have already been made freely available (Canela-Xandri et al., 2017). While Biobanks tend to adopt a cohort design, meta-GWAS studies continue to over-sample cases compared to controls, in order to increase the available power, and are now exceeding 100,000 cases and controls in single studies (Michailidou et al., 2017).

The gold standard for evaluating performance of summary statistic methods is through analysis of simulated data, allowing inference to be compared to a known “truth”. A common method used by GWAS simulators is to procede by adding phenotypes to a sample of genotype data that is either simulated or from a reference population (“forward simulation”). This approach, in particular, can be used very flexibly for generating multiple (quantitative) phenotypes, a design also common to Biobank datasets (Meyer and Birney, 2018). However, this method does not lend itself to simulating case-control data, since it simulates cases in proportion to what we would expect to see in the reference population; typical GWAS designs recruit cases disproportionally to their frequency in the population in order to increase power. In order to forward simulate a GWAS cohort, we would need to simulate until we had the required number of cases and controls, discarding additional samples (typically a large number of controls as cases are normally a minority in the population). This is compuationally expensive, and wasteful.

Instead, when simulating case-control data, we typically simulate or sample genotype data conditional on a supposed distribution of phenotypes. Simulation options in this case are more limited because the problem is mathematically harder. For single causal variant scenarios, resampling from a reference population conditional on allele frequencies at a target variant may be used. For more complicated causal models, involving multiple variants potentially in LD, GWAsimulator (Li and Li, 2008), TriadSim (Shi et al., 2018) or HAPGEN (Su et al., 2011) can very efficiently simulate haplotypes for cases and controls in small genomic regions. In particular, by incorporating mutations and recombinations, HAPGEN can simulate large populations with only a few hundred reference haplotypes. However, the generation of GWAS summary statistics, eg using SNPTEST (Marchini et al., 2007), requires analysis of the individual level data which can be slow, particularly for logistic models which require iterative optimization at each SNP.

The general approach of simulating both genotype and phenotype on an individual level cannot scale well for Biobank-scale or large meta-GWAS situations, because of the number of individuals required. It is also potentially wasteful - the individual level data are not required when the goal is to evaluate methods that work on summary statistics.

Here, we present an alternative approach, which simulates summary statistics directly, without needing to ever generate genotype data. It scales as a function of the number of SNPs, but is constant with regards to the number of samples, thus making it ideally designed for simulation of summary statistics for large case-control studies.

## Results

### Overview of our approach

We first introduce the mathematical calculations which underpin our method. Given a causal model specifying a region of interest, which SNPs in the region are causal, their effects on disease in the form of odds ratios, and reference data on allele and haplotype frequencies in controls, we calculate the expected *Z* score from a Cochran Armitage score test under an additive model at each SNP in the region. (Cochran Armitage score tests have been used for GWAS because of their computational simplicity, requiring no iterative maximisation procedure, and because they allow for additive, dominant or recessive coding, although additive coding is the most commonly used (Sasieni, 1997)).

Simulated Z scores can then be derived by multivariate normal simulation using standard software, with the variance-covariance matrix calculated from correlations between the SNPs in the reference data. This suffices in the case where the summary statistic methods to be used work upon Z scores alone. However, when log odds ratios and their standard errors are required, we appeal to the asymptotic similarity of score tests and Wald tests, and simulate standard errors under the causal model. Together with simulated Z scores, we can then back-calculate the log odds ratios as the product of simulated Z scores and standard errors. An outline description of our calculations follows; full details are given in the Supplementary Information.

Let *Y*_*i*_ ∈ {0, 1} denote the indicator of disease status for the *i*th of *N* individuals sampled according to case-control status (*N*_1_ cases, *Y*_*i*_ = 1; *N*_0_ controls, *Y*_*i*_ = 0). Let *n* be the total number of SNPs. For any SNP *X*, write 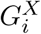 for its genotype coding ∈ {0, 1, 2} at sample *i*. Then, for the commonly used Cochran-Armitage score test, the Z-Score at SNP *X* is computed as:

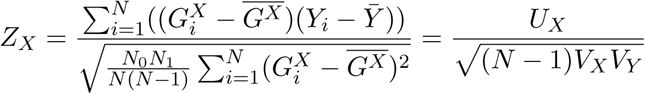

where *V*_*X*_, *V*_*Y*_ denote var(*X*), var(*Y*), respectively.

Write **W** = (*W*_1_*, …, W*_*m*_)^*T*^ for the vector of causal SNPs and *γ* = (*γ*_1_*, …, γ*_*m*_)^*T*^ for their log odds ratios of effect. We assume that *Y*_*i*_ given 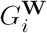 can be modelled as a binomial logistic regression:

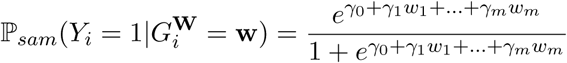

where ℙ_*sam*_() denotes that this is the probability within the GWAS sample and *γ*_0_ is chosen such that 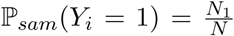. The conditioning is required because allele frequencies vary between cases and controls at causal variants and those in LD with them, meaning the overall allele frequencies in our sample differ from those in the population as a whole. By specifically distinguishing between ℙ_*sam*_() and the more general ℙ(), we can condition on having chosen *N*_0_ controls and *N*_1_ cases and thus perform the conditional simulation needed for case-control studies.

By conditioning upon the values of *G***^W^** and *Y*, we obtain the expected value of *U*_*X*_, the covariance between *G*^*X*^ and *Y*:

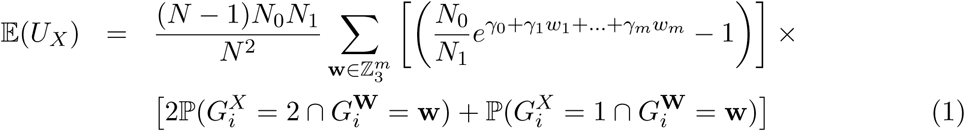

The variance of *U*_*X*_ is *V*_*X*_*V*_*Y*_ where 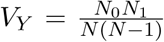 and *V*_*X*_ is the variance of *G*^*X*^. As *V*_*X*_ is a variance, a natural model is an inverse gamma distribution, *V*_*X*_ *∼* Γ^-1^(*α, β*). By similar conditioning upon *G***^W^** and *Y*, we show that the parameters of this distribution are

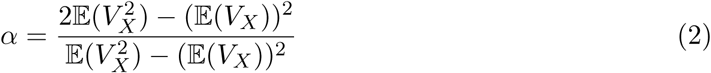

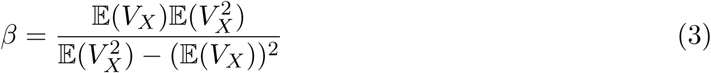

(the derivation of this and expressions for the first two moments of *V*_*X*_ are given in the Supplementary Information). This means we can either simulate *V*_*X*_ from its distribution or calculate

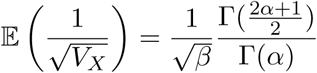

so that, to a first order approximation,

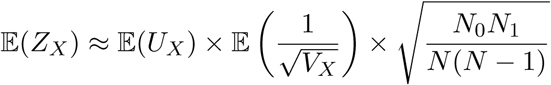

Putting this together, we can now calculate the expected *Z* score, **Z**^E^, across a set of SNPs, given a causal model and some phased reference data with which to calculate the probabilities in (1). Note that the computational complexity of this calculation is independent of both disease frequency and the number of samples required.

For some applications, the expected Z Score may suffice. However, note that the expected GWAS p value is not the p value associated with the expected Z score. Instead, we must simulated “observed” GWAS results which vary randomly about **Z**^E^, with variance 1, such that the correlation between the *Z* score at two SNPs is equal to the correlation between their genotypes (Burren et al., 2014). It is hence computationally simple to simulate multiple realisations of GWAS *Z* scores as **Z**^*^ *∼* MVN(**Z**^E^, **Σ**), where **Σ** is a matrix describing correlation between SNPs for the region, again estimated from the reference panel.

To generate log odds ratios, *γ*, and their standard errors, *σ*, we appeal to the asymptotic similarity of Wald tests from a logistic regression model to the Cochran Armitage score test, and the result that the variance of the score statistic *U*_*X*_ is the inverse of the variance of the estimated *γ*, under the null (McCullagh and Nelder, 1983). Thus, we simulate 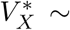 Inverse Gamma(*α, β*) with (*α, β*) given by (2,3) and hence 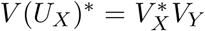. Finally, we set 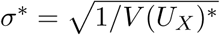 and calculate *γ*^*∗*^ = *σ*^*∗*^**Z**^*^.

### Validation of simulated summary statistics

We visually confirmed that the calculated **Z**^E^ appeared sensible for a selection of one to four independent causal SNP models in a single region (Figure S1). We next simulated data using our method or an individual-based method (HAPGEN+SNPTEST) for five scenarios (Table 1) for a more detailed evaluation. Note in particular the difference between scenarios 4 and 5. In scenario 4, two variants in weak LD each have a log odds ratio of log(1.2) = 0.18 or log(1/1.2) = −0.18. In this case, marginal estimates of odds ratios are close to these values, and Z scores are highly significant. In scenario 5, the pair of odds ratios are the same, but at strongly linked variants (*r*^2^ = 0.8). This would be expected to cause the effect of one to be “cancelled” by the other in the marginal associations, so that estimates of log OR are attenuated towards 1 and significance is dramatically lower, as seen for both HAPGEN+SNPTEST and simGWAS simulations (Figure S2). Visually, the Manhattan plots generated by the two methods were similar (Figure 1). However, we did notice that simGWAS displayed greater variability than HapGen+SNPTEST at the SNPs with smallest p values. Formal comparison of the distribution of statistics showed that the mean log OR and mean Z score were statistically indistinguishable between the two methods, but that simGWAS produced results with greater variability, resulting in some statistically significant differences in Kolmogorov-Smirnov tests comparing the two distributions (Table S1).

**Table 1:**
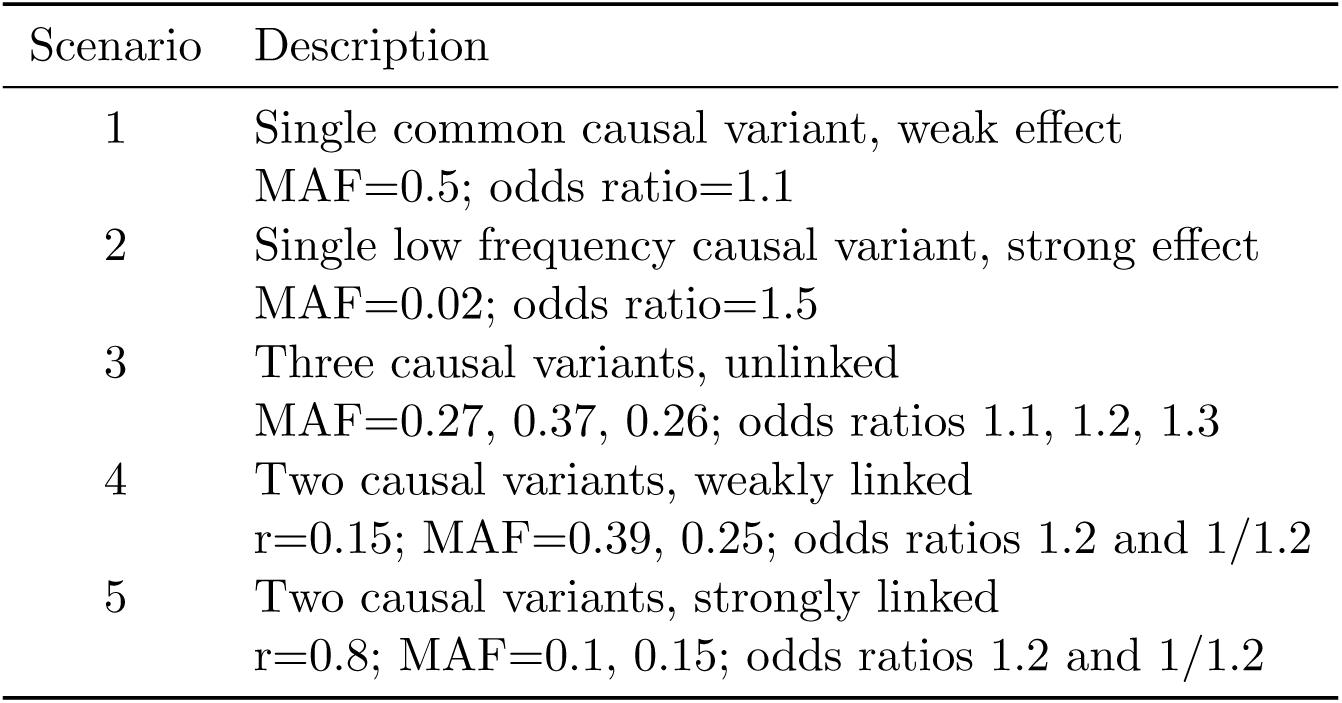
Five simulation scenarios considered for validation of results

| Scenario | Description |
| --- | --- |
| 1 | Single common causal variant, weak effect<br>MAF=0.5; odds ratio=1.1 |
| 2 | Single low frequency causal variant, strong effect<br>MAF=0.02; odds ratio=1.5 |
| 3 | Three causal variants, unlinked<br>MAF=0.27, 0.37, 0.26; odds ratios 1.1, 1.2, 1.3 |
| 4 | Two causal variants, weakly linked<br>$r=0.15$; MAF=0.39, 0.25; odds ratios 1.2 and 1/1.2 |
| 5 | Two causal variants, strongly linked<br>$r=0.8$; MAF=0.1, 0.15; odds ratios 1.2 and 1/1.2 |

**Figure 1:**
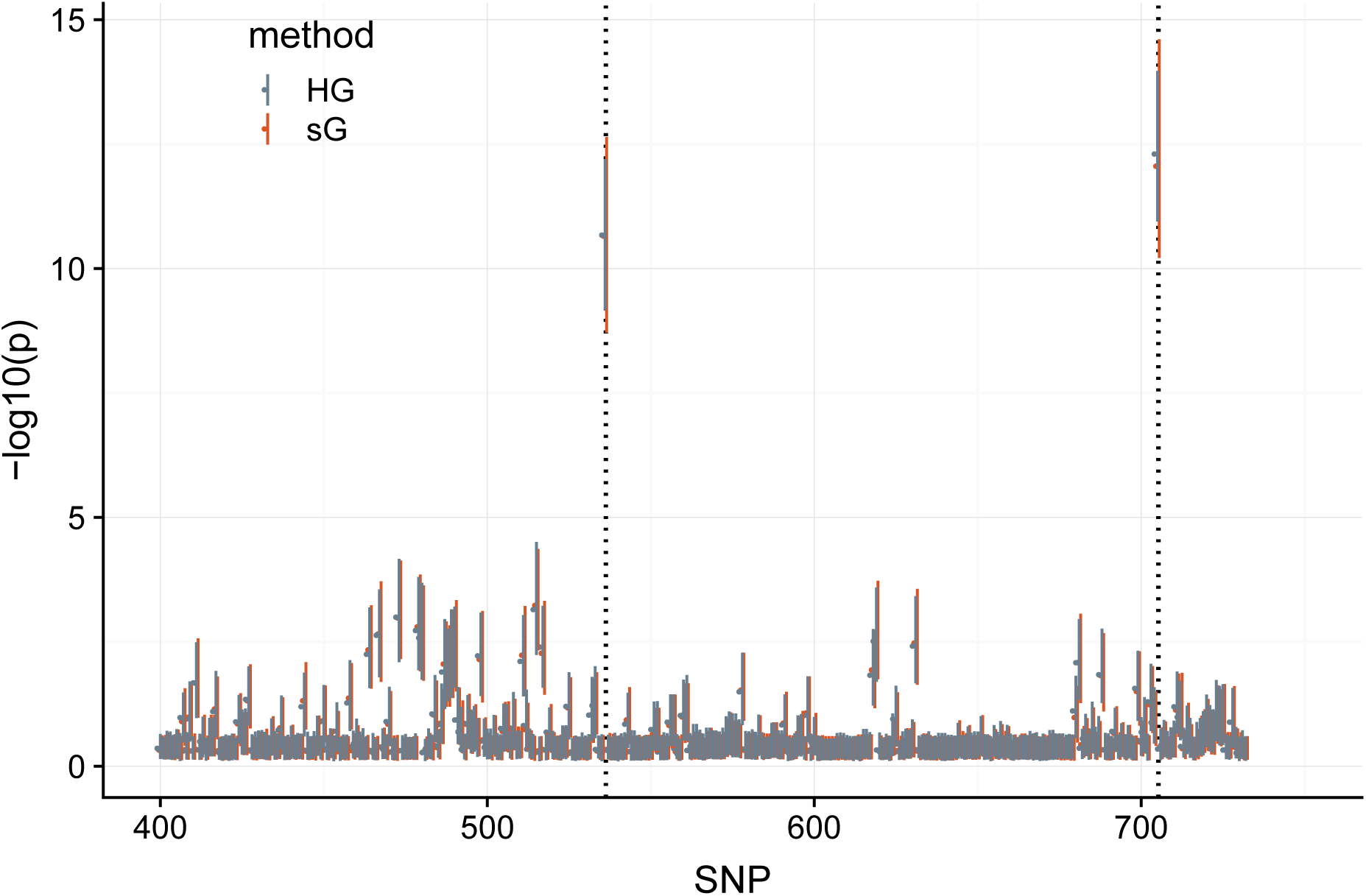
Results from simGWAS (sG) are visually similar to those from HAP-GEN+SNPTEST (HG). The figure shows *∼*350 SNPs from around the causal variants in the simulated region under scenario 4, with 5000 cases and 5000 controls. Points show the median -log10(p value) for each SNP, and ranges the IQR across 1000 simulations. Location of causal variants are marked with dotted lines.

To investigate this, we conducted forward simulations at the causal variants only in each scenario, as a gold standard, and found that results from simGWAS more closely matched those from this gold standard than did those from HapGen+SNPTEST (Figure 2, Table S1).

**Figure 2:**
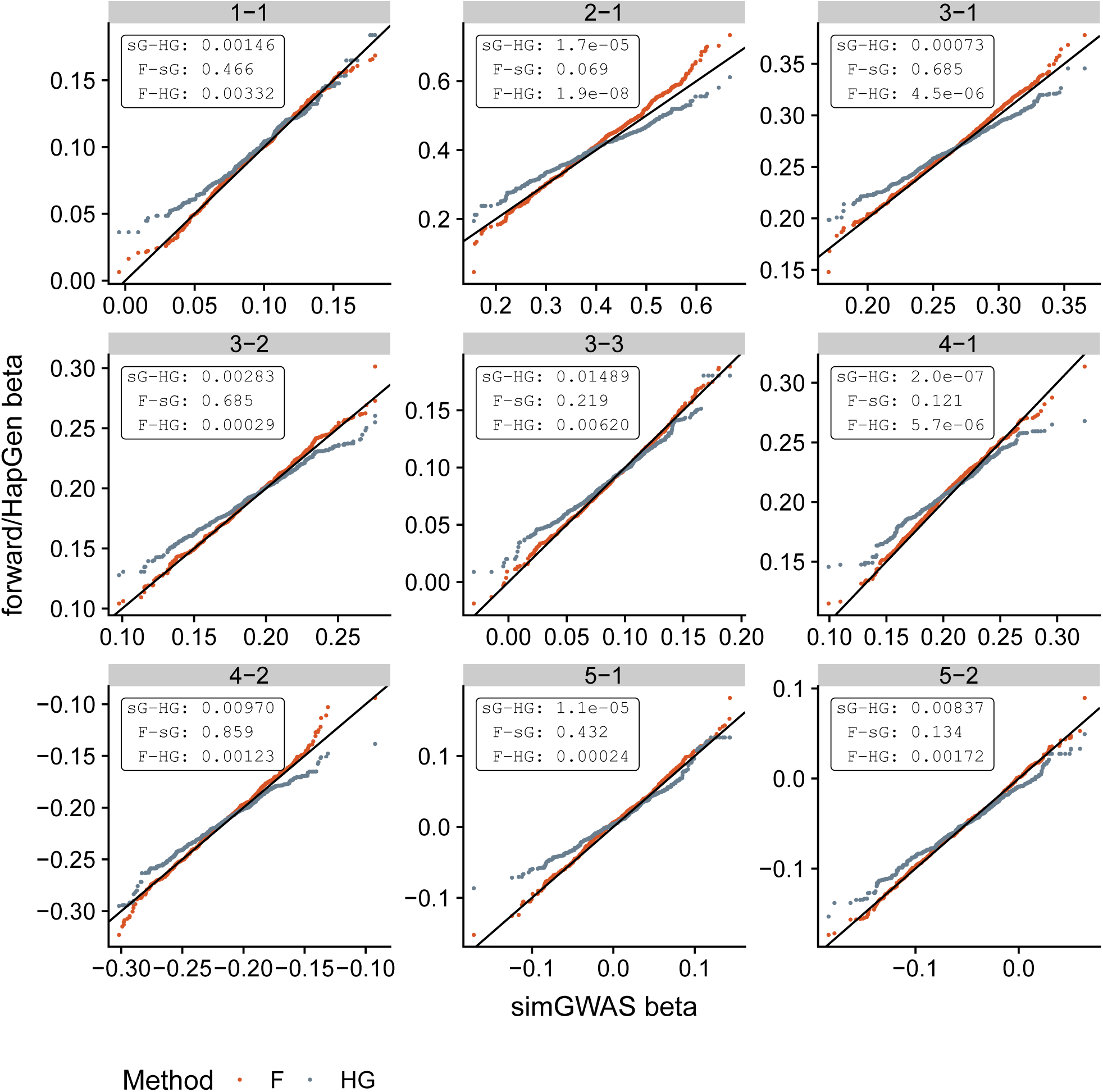
QQ plots comparing distribution of log OR at causal SNPs across 1000 simulations with 5000 cases and 5000 controls. Each plot compares the distribution of log OR generated by simGWAS (sG, x-axis) to that from HapGen+SNPTEST (HG) or forward simulation (F). Distributions were compared using Kolmogorov-Smirnov, and p values are shown in the top-right of each subplot. The label of each plot gives the corresponding “scenario-snp” pair - i.e. the label 3-1 refers to scenario 3, first causal SNP.

Finally, we compared simulation speed of each strategy as the number of causal variants, the number of samples and the number of replicates varied. For a region with 1000 SNPs using AFR data from 1000 Genomes (~ 600 samples), both methods were very fast (< 30 seconds) for the simplest scenario of 1000 cases and 1000 controls. We found that both methods required slightly, but negligibly, more time as the number of causal variants increased from one to six (Figure 3a). As expected, HAPGEN+SNPTEST scaled linearly with either the number of replications (number of complete sets of data simulated from the same scenarios) or sample size, whereas simGWAS timings were independent of either factor (Figure 3b–c). This emphasizes the potential for fast simulation of summary statistics for very large case-control datasets.

**Figure 3:**
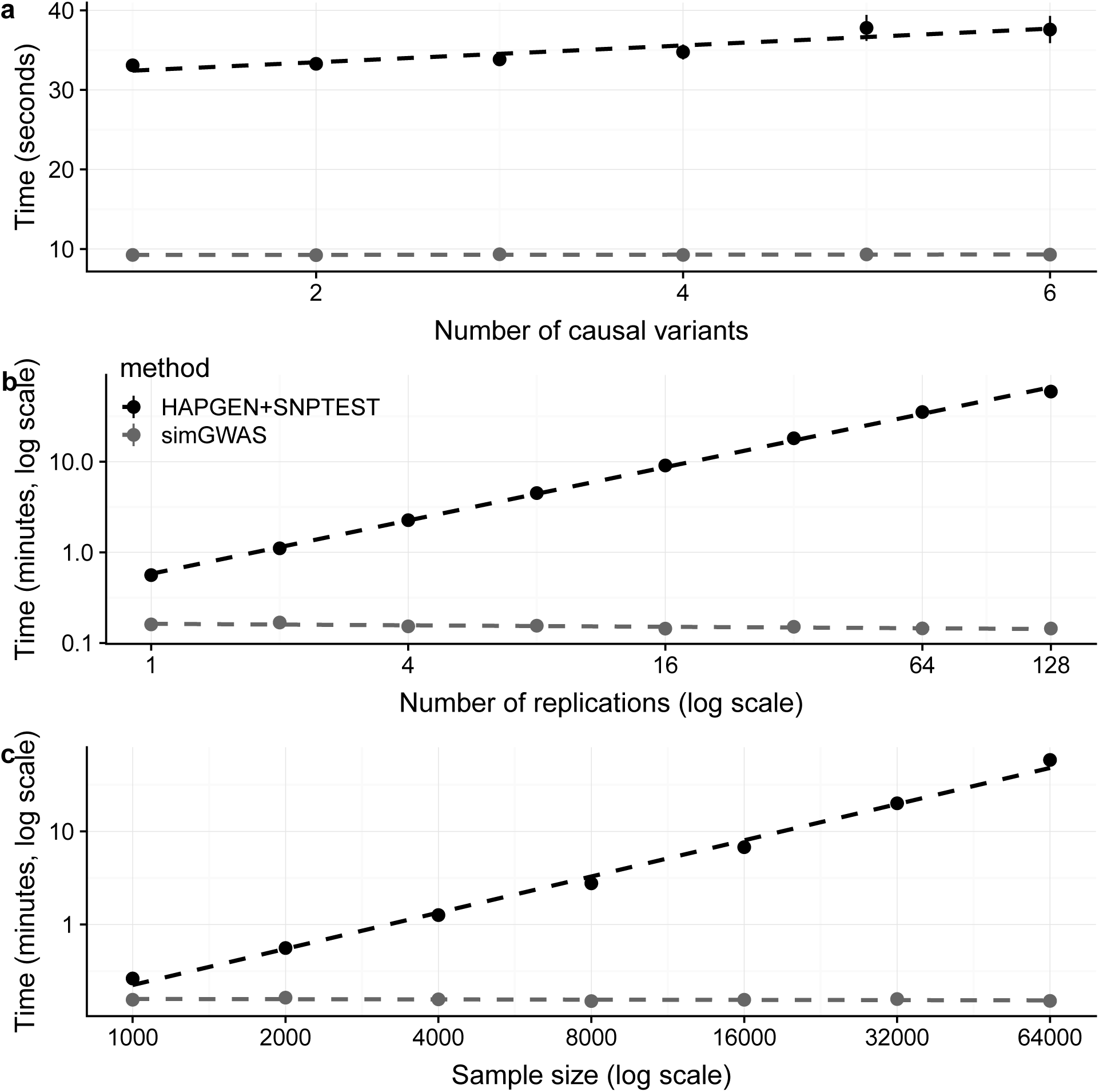
Time taken to perform simulations under simGWAS or HAPGEN2+SNPTEST2 strategies. simGWAS is given in brown and HAPGEN2+SNPTEST2 in blue. Each point represents the mean time for 50 independent runs of the software, with standard deviation about that mean indicated by the vertical bars. **(a)** The effect of number of causal variants on run time. 2000 cases, 2000 controls, single replication, causal variants varying from 1 to 6. **(b)** The effect of number of replications on run time. 2000 cases, 2000 controls, 2 causal variants, number of replications varying from 1 to 100. **(c)** The effect of sample size on run time. single replication, 3 causal variants, number of cases and controls (each) varying from 1000 to 64000.

## Discussion

Simulating GWAS summary statistics in the context of case-control studies, for any required causal model and set of odds ratios, has several potential applications. Primarily, simulated GWAS results have become the accepted gold standard for validating newly developed statistical models for the analysis of GWAS data. Our intent is to enable the faster simulation of summary statistics compared to individual level data simulation, while at the same time using considerably less disk space. Although the method focuses on region-level simulation, it can be used to generate genomewide statistics if required, by breaking the each chromosome into approximately independent blocks, according to recombination hotspots or breakpoints derived from examining correlation between genotypes Berisa and Pickrell (2016).

We note that our method depends on an assumption of additivity - across alleles at each SNP, and across causal SNPs in a region. This additive model is used by the overwhelming majority of GWAS analyses, by genetic risk score approaches, and by LD regression methods (Dudbridge et al., 2018; Bulik-Sullivan et al., 2015). Thus it seemed the sensible place to start. However, examining how these methods which assume additivity perform when the underlying model is *not* additive is an interesting research question. A future direction to extend our method could be to adapt it to simulate data under any genetic model by expressing disease risk as a function of genotypes at causal SNP haplotypes. While we have focused on retrospective case-control designs as an obvious gap in the GWAS simulation toolbox, our methodology could be relevant in the area of extreme-sampling designs, where power is maximised for a fixed cost by sampling individuals with extreme values of a quantitative trait, for example in a study of blood pressure(Warren et al., 2017). We could adapt our method to this design by expressing the distribution of haplotype frequencies as a function of a quantitative trait.

In addition to supporting method development, simulation of GWAS statistics is also used in tests aggregating information across sets of SNPs, e.g. for pathway analysis. Pathway analysis can test either the *global null*, of no association between any SNP and phenotype, or the *competitive null*, which assumes there are some truly associated SNPs, but that these are randomly distributed amongst the sets of SNPs considered (i.e. those near genes in or out of the pathway under test, or those corresponding to presence or absence of a feature of interest). The second seems more appropriate, because it acknowledges that enrichment tests are performed in the context of genome-wide significant associations having been already found. However, the second is also much harder to simulate.

A common technique for simulating under a competitive null is permutation testing; the underlying dataset is maintained, and labels are permuted to generate new datasets where traits are still associated, but there is no possible correlation to the feature of interest. However, doing this so as not to destroy the genomic structure within the region, can require inventive generation of null distributions, for example, by circularisation and permutation of genomic features to allow empirical null distributions to be calculated under a competitive null (Trynka et al., 2015). While these are efficient, they can only be used for features that span shorter distances than LD - e.g. for chromatin mark enrichment but not genes collected in pathways.

To allow more simple simulation techniques to be used, pathway-based tests of the competitive null have been adapted to have the same expected null distribution as tests of the global null. This requires replacing p values for individual genes by their ranks (Evangelou et al., 2012) which loses distributional information.

There is therefore potential to further develop pathway or enrichment test methodology if the distribution of test statistics under a competitive null hypothesis could be derived. Our method would naturally allow simulation of GWAS summary data under a specific hypothesis about the location and magnitude of genetic effects, in order to generate empirical null distributions for tests of the competitive null, preserving genomic structure even when analysis is performed across multiple regions.

Finally, our method could be used to evaluate output of fine-mapping applied to real data. Particularly in regions where the patterns of LD between putative associated SNPs are complex, it can be hard to dissect what the true causal variants are. Different fine mapping methods make different assumptions about the number and independence of causal variants, which can produce conflicting results (Wallace et al., 2015; Newcombe et al., 2016). By generating expected summary statistics under alternative fine-mapped solutions, it may be possible to see whether one or another is more compatible with observed data.

Our method enables faster simulation of GWAS case-control summary statistics compared to individual level data simulation, at the same time using considerably less disk space. This should facilitate computationally simpler evaluation of existing and new summary GWAS methods and has the potential to underpin new method development in other areas.

## Methods

### Simulations to validate summary statistics

We evaluated our proposed method by simulating summary statistics in parallel using simGWAS (our method) and the same settings with HAPGEN2 + SNPTEST2, using reference data from 1000 Genomes Phase 3 (1000 Genomes Project Consortium et al., 2015) (AFR cohort, ~ 600 subjects). Reference data was downloaded from https://mathgen.stats.ox.ac.uk/impute/impute_v2.html#reference. We visually compared distributions of summary statistics, as well as time to create the statistics under different scenarios. Full code to run these simulations is available from http://github.com/chr1swallace/simgwas-paper.

### Funding

MF and CW are funded by the Wellcome Trust (WT099772, WT107881) andCW by the MRC (MC UU 00002/4). MF is currently funded by Dementia Platforms UK.

### Availability

A software package making our method available is at http://github.com/chr1swallace/simGWAS and code used to produce the figures and results in this paper is available at http://github.com/chr1swallace/simgwas-paper.

### Conflict of interest

None to declare

## Acknowledgments

We thank Marcus Klarqvist for advice on efficient C coding.

## Supplementary Information

### Cochran-Armitage test of association

For a GWAS dataset, let *Y*_*i*_ ∈ {0, 1} denote the indicator of disease status at the *i*th sample. Let there be a total of *N* samples selected, with *N*_1_ having been chosen from disease cases (*Y*_*i*_ = 1) and *N*_0_ having been chosen from disease controls (*Y*_*i*_ = 0). Since this sampling is conditional upon case/control status, genotype frequencies may differ between our *N* samples and the whole population at disease associated SNPs. We therefore need to distinguish between which datasets the genotype probabilities are from; write ℙ_*sam*_ for probabilities computed for the samples (i.e. 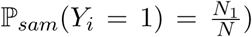, and ℙ for probabilities generated with reference to the whole population.

Let *n* be the total number of SNPs. For any SNP *X*, write 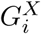 for its genotype coding ∈ {0, 1, 2} at sample *i*.

For the commonly used Cochran-Armitage test, the Z-Score at SNP *X* is computed as:

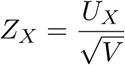

Where:

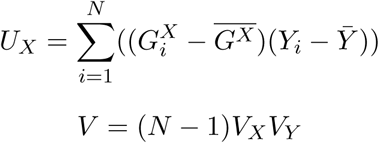

and *V*_*X*_, *V*_*Y*_ are the variance of *G*^*X*^ and *Y* respectively:

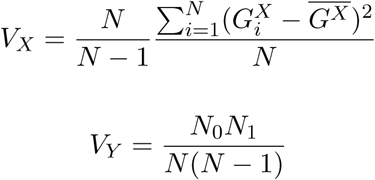

i.e.:

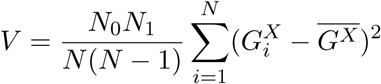

Under the null hypothesis of no association at SNP *X*, *Z*_*X*_ is distributed as a standard normal. Hence the two-sided p-value at *X* is given by:

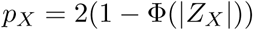

where Φ is the cumulative distribution function of the standard normal distribution. Conversely, given the unsigned p-value at *X*, the absolute value of the Z-Score is:

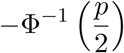

### Allelic frequencies under a Causal Model

Write **W** = *W*_1_*, …, W*_*m*_ for the vector of causal SNPs. From phased publicly available reference datasets such as UK10K (Walter et al., 2015), it is possible to estimate haplotype frequencies across all SNPs in **W** at any subset of potential causal SNPs in control datasets. Since they are causal, these frequencies will differ in cases, and it is those frequencies we derive first. Note that, since sampling dependent only upon case/control status, we can assume:

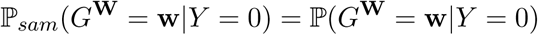

Write *γ*_1_*, …, γ*_*m*_ for the log odds ratios of effect for the causal SNPs in the population. We assume that *Y* given *G***^W^** can be modelled as a binomial logistic regression. Then, from (Prentice and Pyke, 1979), the sample-specific odds ratios are the same as those at the population-level, and we can write:

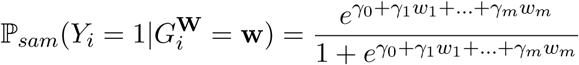

where *γ*_0_ is an intercept parameter. Since GWAS sampling is retrospective, the proportion of cases in the sample is fixed at 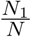, constraining *γ*_0_, which can be computed as follows:

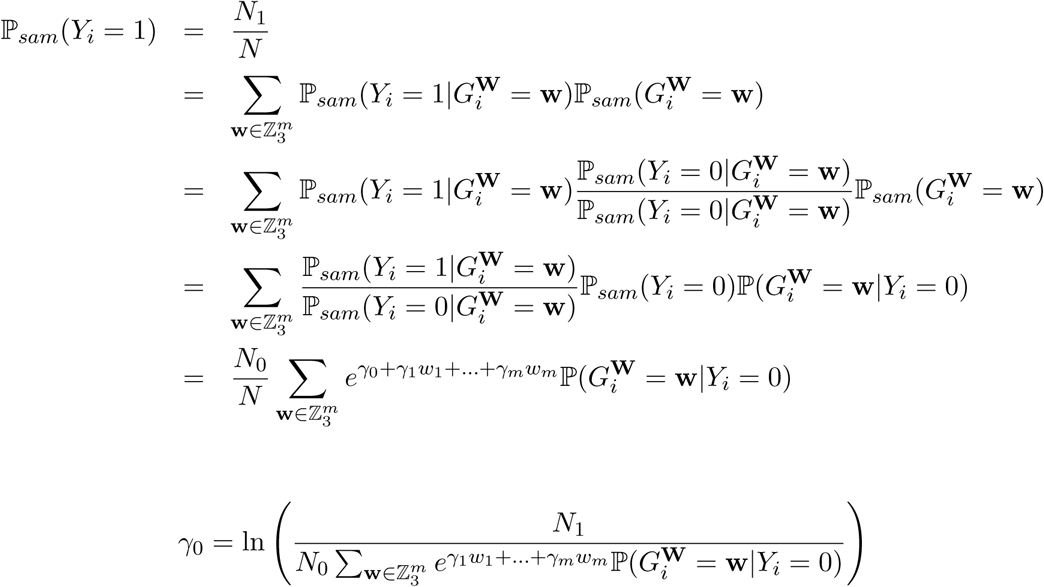

Hence we can compute:

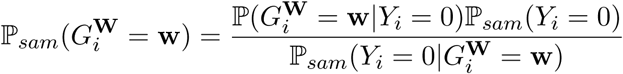

And also:

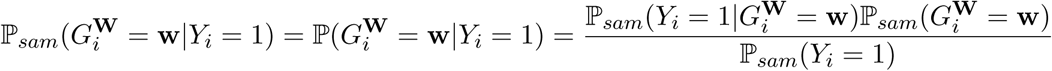

To derive genotype probabilities at SNPs in LD with the causal SNPs, we assume that LD structures do not differ between cases and controls, and hence the correlation between **W** and *X* is independent of both disease status and our sampling. Thus:

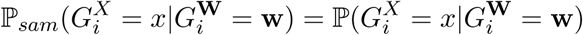

and we can estimate, for both the whole population, and for our sample:

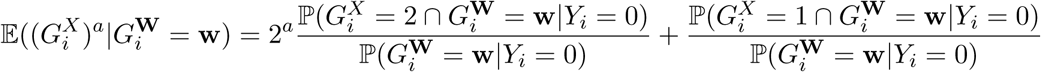

from our reference dataset, for any constant *a*. From this, we compute:

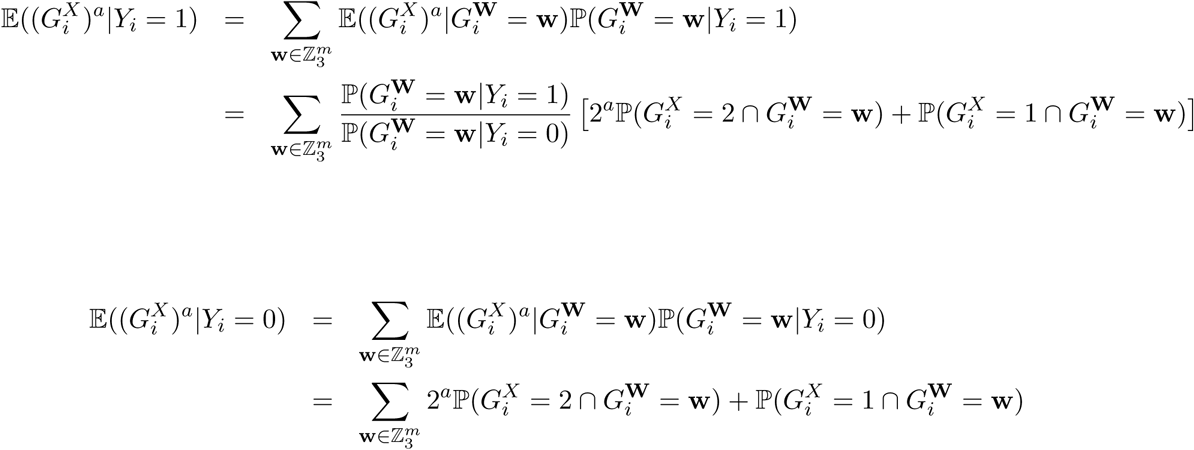

By expanding out the numerator in terms of probabilities within the sample dataset, we see that:

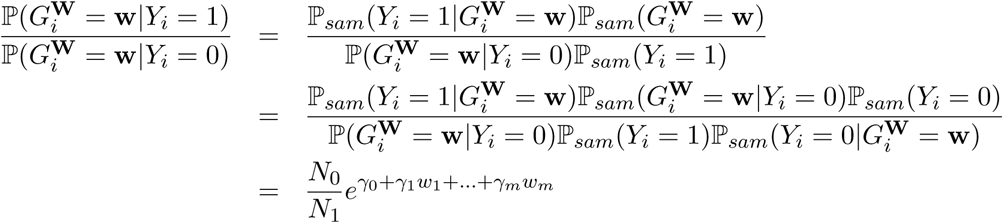

And hence:

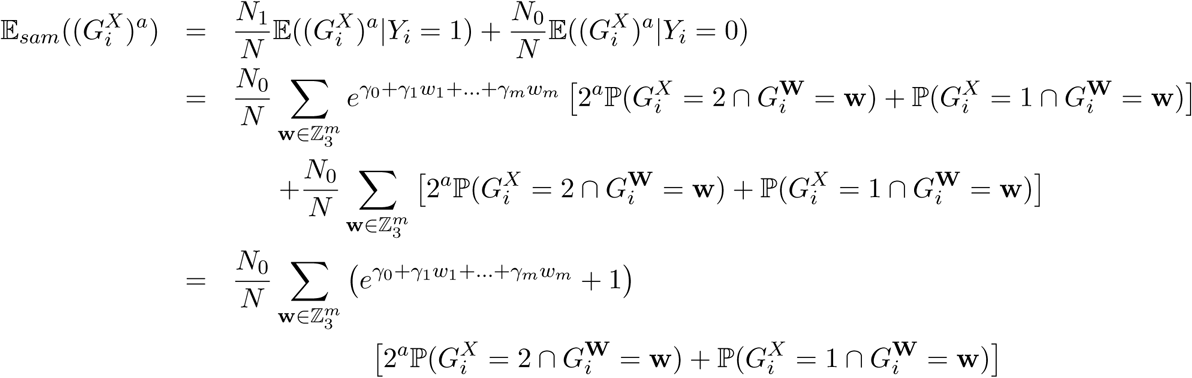

### Estimation of Z Score for the causal model given by W and *γ*

Finding the true expectation of 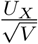 is intractable, so instead we compute a first order approximation by assuming independence:

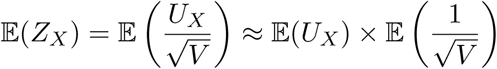

These terms can be computed as shown in the following sections.

#### Estimation of *U*_*X*_, the covariance between *G*^*X*^ and *Y*, for the causal model given by W and *γ*

We compute the expectation of *U*_*X*_ in our sample as follows:

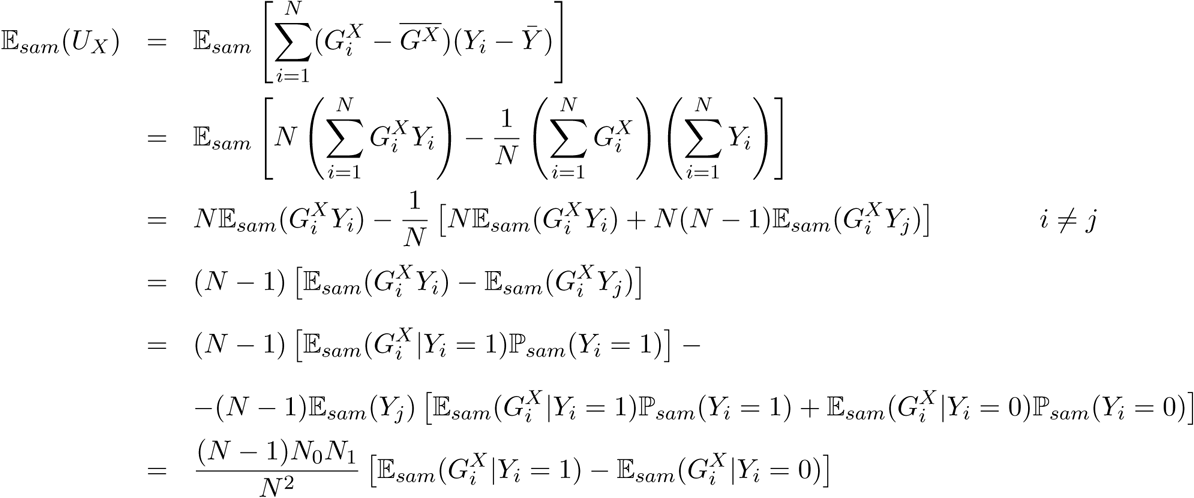

Using the expressions for 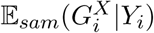 given in Section, this becomes:

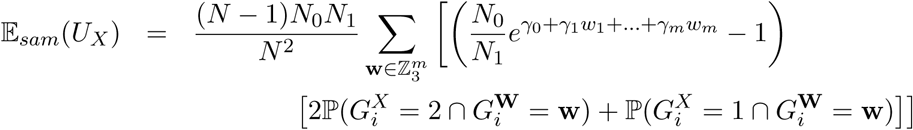

#### Estimation of *V*_*X*_, the variance of *G*^*X*^, for the causal model given by W and *γ*

Recall:

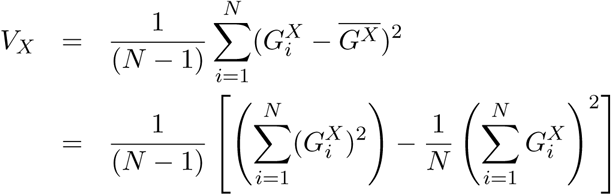

This is tractable, however, we need to find 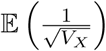, which is more complex.

*V*_*X*_ is the variance of a normal, and so we model it as an Inverse Gamma (*α, β*) distribution. Then 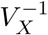 has a Γ(*α, β*^-1^) distribution, and 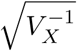 has a generalised gamma distribution with parameters 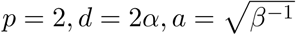. If *V*_*X*_ ~ Inverse Gamma (*α, β*), then

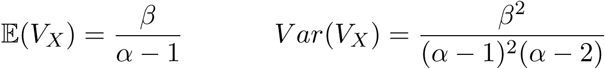

Assuming we have computed 𝔼 _*sam*_(*V*_*X*_) and 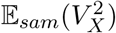, *α* and *β* are completely specified as:

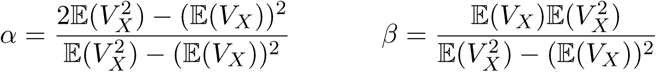

and 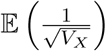 may be simply computed using:

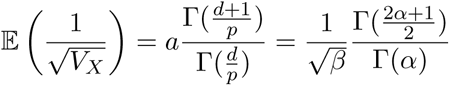

#### Expectation of *V*_*X*_

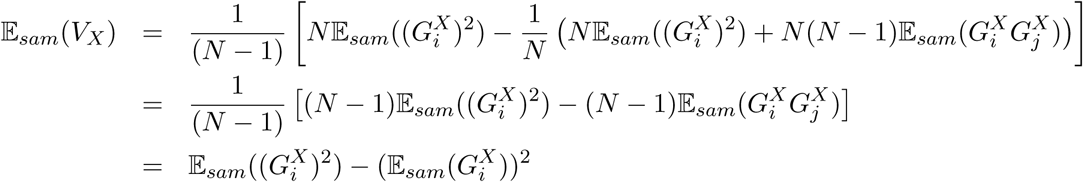

#### Expectation of 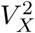

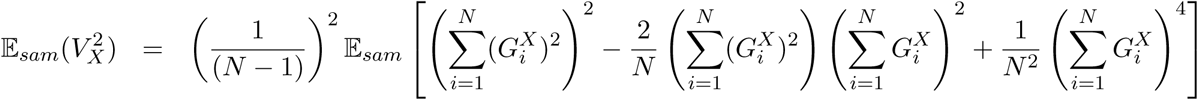

Let 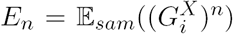. Breaking this down into terms, for (*i, j, k, l*) representing different indices, we have:

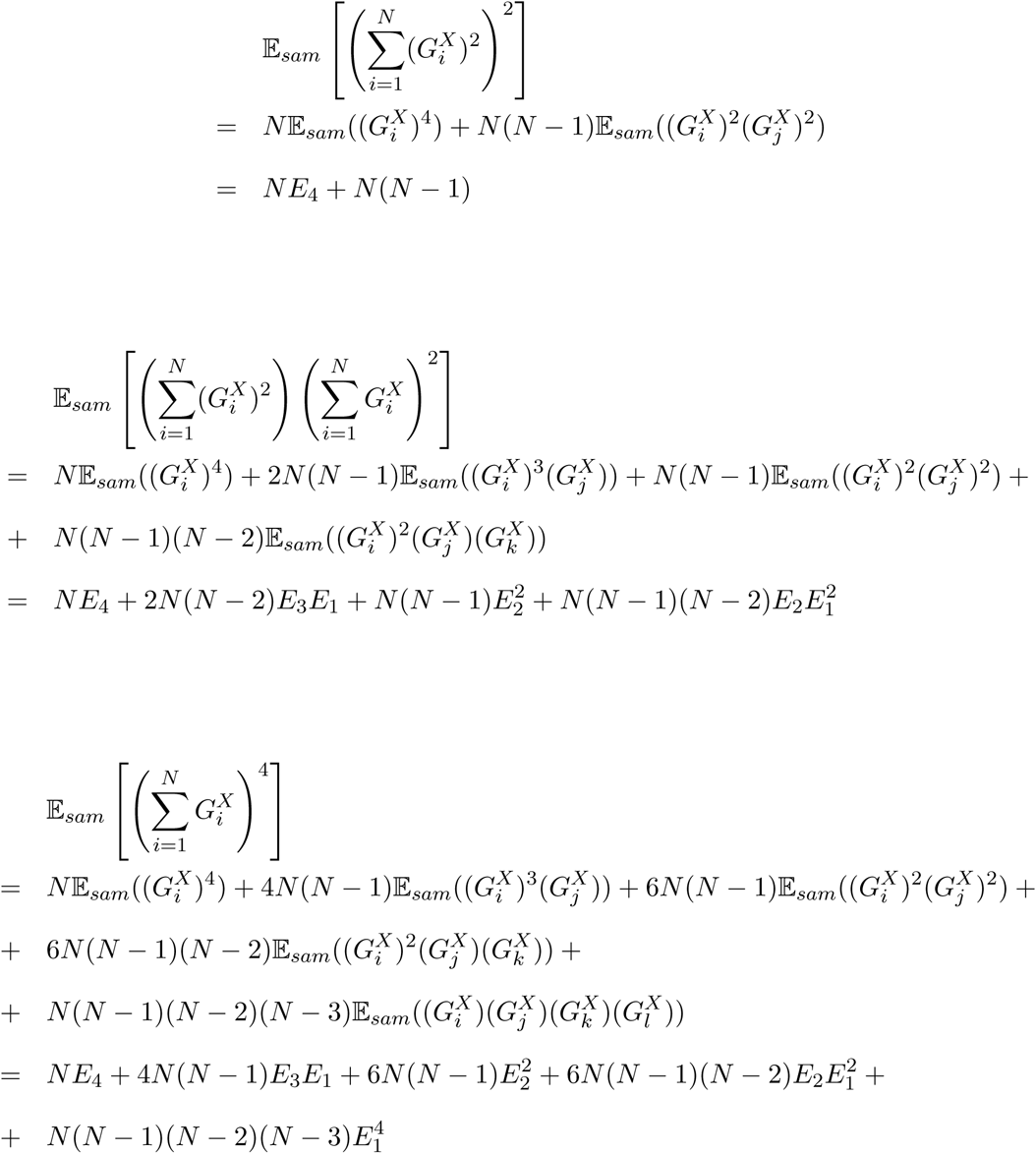

Giving:

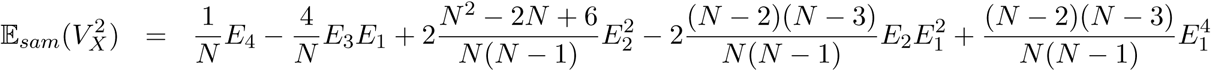

### Summary

Thus, given only a choice of which SNPs are causal (**W**), their effect sizes (***γ***), sample sizes (*N*_0_*, N*_1_) and a reference dataset from which we can derive allele frequencies 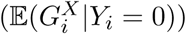 and the relationships between SNPs 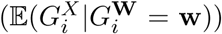, we can derive an expected Z Score, **Z**^*EXP*^ at any SNP, causal or not.

This can then be used directly. However, most applications require simulated output from such a GWAS. **Z**^*SIM*^ can therefore be computed, which will be distributed:

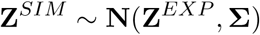

where **Σ** is the genotype correlation matrix for the SNPs in this region (Burren et al., 2014).

Table S1 (file: simgwas-supptab.csv). Comparison of beta (log OR) and Z scores from simulations run with simGWAS, HapGen, and forward simulation. Mean and standard deviations are given at causal variants and one unlinked variant per scenario. Means were compared using T tests with Welch extension to accommodate unequal standard deviations, and p values are shown in the columns “.meantest”. Formal comparison of the full distribution was conducted by Kolmogorov-Smirnov tests (KS) and p values are shown in the columns “.KStest”. The sample size (n) is the number of cases and controls - i.e. n=1000 indicates the simulations related to 1000 cases and 1000 controls. Each plot summarises 1000 simulations. The scenario label gives the corresponding “scenario-snp” pair - i.e. the label 3-1 refers to scenario 3, first causal SNP. Unlinked variants are denoted as SNP 0.

---

**Figure S1:**
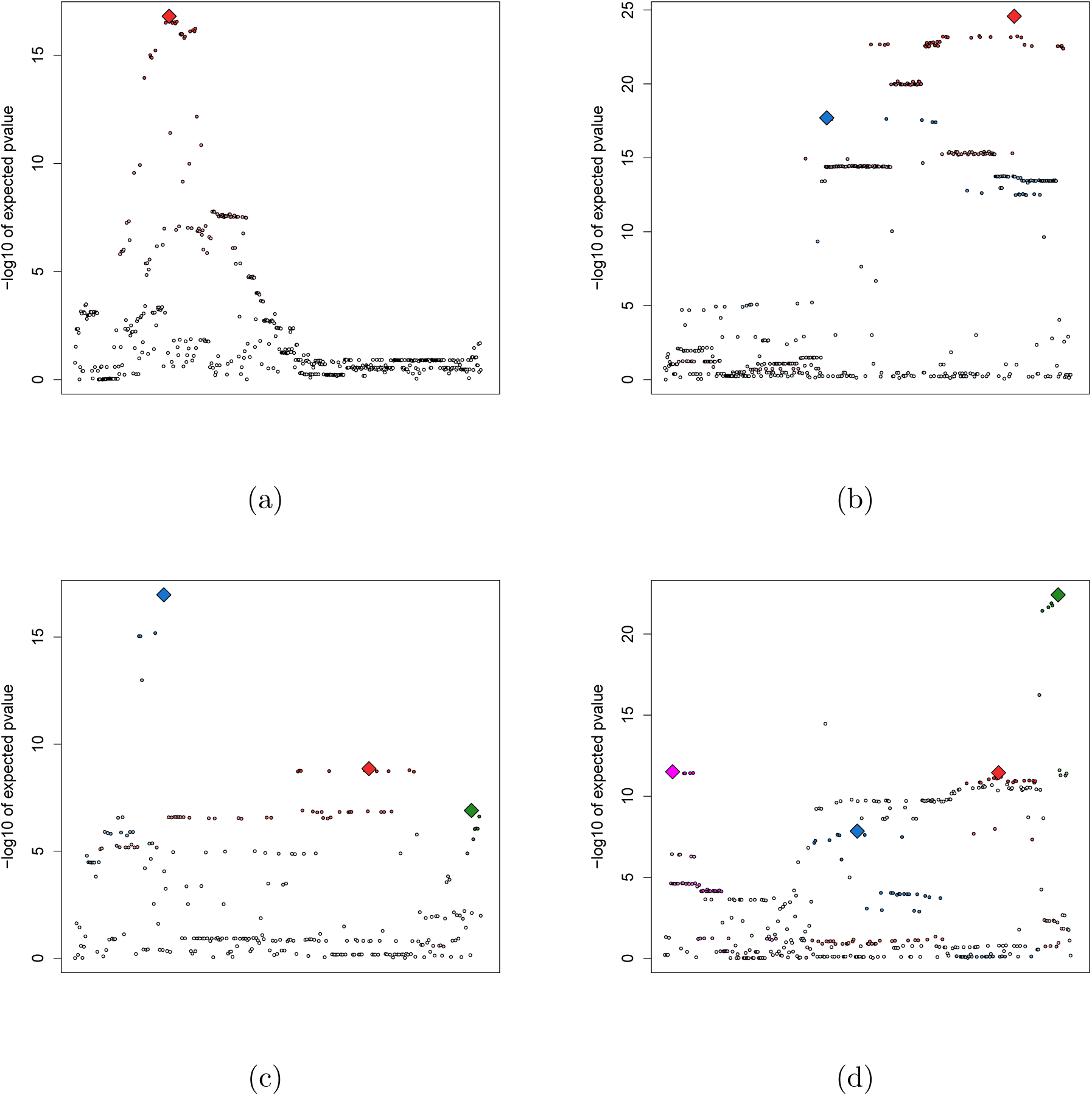
Local Manhattan plots for *p* values generated from expected *Z* scores under different scenarios, in order to confirm by visual inspection that the expected Z Scores produced by our algorithm are consistent with the behaviour we would expect from their causal SNPs. In order to easily see the pattern of association, causal variants chosen were common, with a strong effect and (in the case of multiple causal variants) only weakly linked. Causal SNPs are designated by a coloured diamond. Non-causal SNPs are designated by a circle, coloured according to their LD with their most correlated causal SNP. In each scenario, 5000 cases and 5000 controls were simulated. (a) A single causal variant with MAF = 0.34 and Odds Ratio of effect = 1.3 (b) Two causal variants with MAF = (0.14, 0.30) and Odds Ratio of effect = (1.5, 1.2) (c) Three causal variants with MAF = (0.12, 0.43, 0.17) and Odds Ratio of effect = (1.2, 1.2, 1.2) (d) Four causal variants with MAF = (0.33, 0.44, 0.17, 0.28) and Odds Ratio of effect = (1.5, 1.5, 1.5, 1.5)

**Figure S2:**
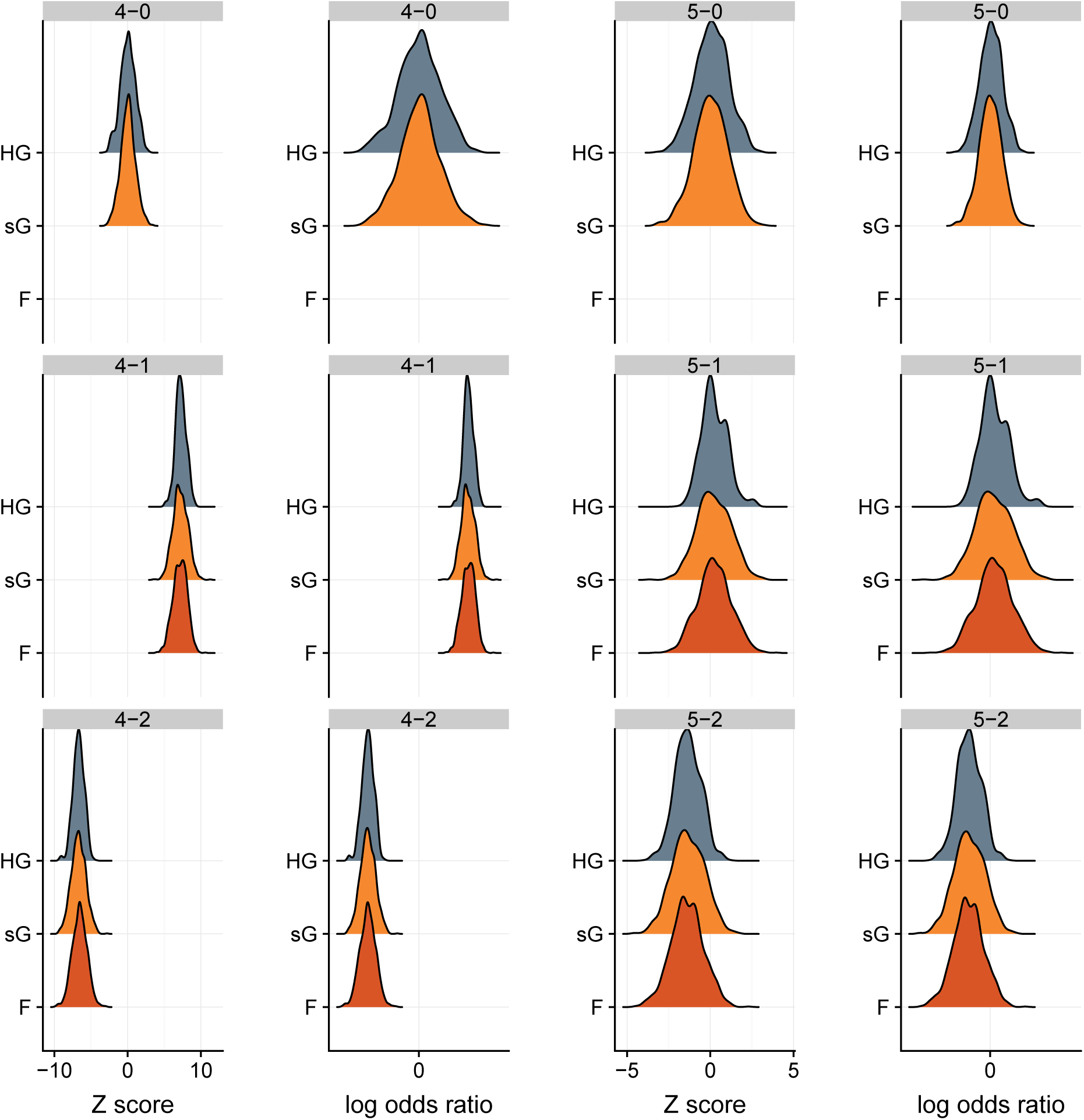
Results from simGWAS (sG) are similar to those from HAPGEN+SNPTEST (HG) and forward simulation (F). Distributions of simulated Z scores and log odds ratios are shown from 1000 simulations assuming 5000 cases and 5000 controls under two scenarios - 4 and 5 - described in Table 1. The label of each plot gives the corresponding scenario-SNP pair, with SNP 0 unlinked (*r*^2^ < 0.15) to either causal variants (no forward simulation results for SNP 0). In scenarios 4 and 5, two SNPs have the same effect sizes but are either weakly linked (*r* = 0.15, scenario 4) or in strong LD (*r* = 0.8, scenario 5). Note that the marginal effect sizes are closer to 0 in scenario 5 because the linked effects cancel when considering marginal association.

